# A Synthetic Single-Stranded Dual-Template Oligonucleotide as a Reference Standard for Monochrome Multiplex qPCR Measurements of Average Telomere Length

**DOI:** 10.1101/2020.06.24.169797

**Authors:** Richard M. Cawthon

## Abstract

Quantitative PCR is frequently used to measure average telomere length (TL) relative to the TL of a reference DNA sample of the investigator’s choosing. This makes comparisons of TLs across studies and laboratories difficult. Here we demonstrate that a single synthetic single-stranded dual-template oligonucleotide (DTO) containing both a telomere repeat sequence (T) and a segment of the human beta-globin (HBB) single copy gene (S) can be used as a universal reference standard for monochrome multiplex quantitative PCR (MMqPCR) measurements of average TL using SYBR Green I as the only fluorescent reporter dye. A set of twelve concentrations of the DTO is prepared by serial 3-fold dilutions, to a lowest concentration of ~20 copies per μl. The 5 highest concentrations are used for the T standard curve, and the 5 lowest concentrations are used for the S standard curve. For each reaction 5 μl containing approximately 3 ng of genomic DNA (or one of the DTO dilutions) is mixed with 5 μl of a 2x MasterMix containing the primers for T and S amplification, and MMqPCR is performed. The design of the primers and thermal cycling profile allows all T amplification signals to be collected before exponential amplification of the S signal begins. Exponential amplification from S is then carried out in a temperature range that keeps the telomere product fully melted and therefore unable to influence the S amplification signal. The T value for each DNA sample is the Standard Curve DTO dilution that contains the same number of copies of the telomere sequence as the experimental sample, and the S value is the DTO dilution that contains the same number of copies of the single copy gene sequence as the experimental sample. Dividing the first dilution by the second dilution yields an absolute T/S ratio, since it is expressed relative to the fixed 1:1 T/S ratio that is built into the DTO by design. Absolute T/S ratios for average TL in 48 human DNA samples determined by this method correlated strongly with mean Terminal Restriction Fragment (mTRF) lengths for the same DNA samples determined by the Southern Blot method (R-squared = 0.801). This DTO and the accompanying protocol may facilitate the standardization of average telomere length measurements and analyses across laboratories.

## INTRODUCTION

Measurement of average telomere length by absolute quantitative PCR using two synthetic oligonucleotides as the reference standards, one for the telomere sequence and the other for the single copy gene (scg) sequence, has been published [1]. This method requires frequent, careful preparation of telomere and scg oligomer solutions with precisely known concentrations.

To simplify absolute qPCR measurements of average telomere length, we here present a dual-template oligonucleotide (DTO) consisting of a telomere sequence (T) and an scg sequence (S) in a single molecule, a design that guarantees the two templates will be present in a 1:1 ratio each time a solution containing them is prepared. To set up Standard Curves for the determination of T and S in an experimental DNA sample, several concentrations of the DTO are generated by serial dilution. The high range of DTO concentrations is then used to determine the T values, and the low range to determine the S values. The absolute T/S for a given experimental DNA sample is then the ratio of the two dilutions of the DTO that match the experimental sample for copy number of the T and S sequences, respectively.

To obtain absolute T/S ratios from singleplex qPCR assays of telomere length, the high DTO concentrations in the T Standard Curve and all the research subjects’ DNA samples can be qPCR-amplified with only the telomere primer pair present; and separately, the low DTO concentrations in the S Standard Curve and all the subjects’ DNAs can be qPCR-amplified with only the single copy gene primer pair present.

Here we present a monochrome multiplex qPCR assay for absolute T/S ratios using this DTO as the reference standard. The advantages of multiplex over singleplex designs in qPCR are improved accuracy, higher throughput, and lower costs. In this monochrome multiplex method, the thermal cycling profile is designed to pause the exponential amplification of the S signal until after all T amplification signals have been collected, as described in a recent study of relative TLs measured by a modified MMqPCR method [2]. During telomere amplification the highest temperature reached in each cycle is high enough to melt the telomere PCR product (amplicon), but too low to melt the GC-clamped beta-globin (HBB) amplicon, thereby putting HBB exponential amplification on hold. After collection of the T signals is complete, high temperature thermal cycling is performed to exponentially amplify and acquire signal from the HBB target, without interference from the fully melted telomere products.

We also show that T/S ratios determined by this new method correlate strongly with mean Terminal Restriction Fragment lengths measured by the traditional Southern Blot method, and decline with age at blood draw across the lifespan, as expected.

## MATERIALS AND METHODS

### Research subjects

The research subjects were 48 individuals (29 females and 19 males, age range 5–94 years) from the Utah CEPH (Centre pour les Etudes du Polymorphisme Humaine) families, families whose participation in medical research contributed to the worldwide effort that built the first comprehensive human genetic linkage maps [3,4]. In the early 1980s bloods were drawn from these subjects, and genomic DNA was extracted using standard phenol-chloroform-based procedures and stored long-term in TE buffer (10mM Tris–HCl, 0.1mM EDTA, pH7.5) at 4°C at a concentration of ~100 ng/ul. This study was approved by the University of Utah’s Institutional Review Board and by the university’s Resource for Genetic and Epidemiologic Research. All participating families provided signed, paper-based informed consent.

### Determination of mean TRF length

Southern blot measurements of mean Terminal Restriction Fragment (mTRF) lengths in these samples has been described previously [5].

### Primer and dual-template oligonucleotide sequences

The telomere and beta-globin (HBB) primer sequences used for this study were identical to those previously reported [6]:

~~~
telg: 5’-ACACTAAggtttgggtttgggtttgggtttgggttagtgt-3’
telc: 5’-TGTTAGGtatccctatccctatccctatccctatccctaaca-3’
HBBu: 5’-CGGCGGCGGGCGGCGCGGGCTGGGCGGcttcatccacgttcaccttg-3’
HBBd: 5’-GCCCGGCCCGCCGCGCCCGTCCCGCCGgaggagaagtctgccgtt-3’
~~~

The single-stranded dual-template oligonucleotide (DTO), ordered from Integrated DNA Technologies, Inc., contains 52 bases of HBB sequence, followed by a C9 linker (iSp9), followed by 35 bases of telomere sequence, ending with a 3’ terminal phosphate to block the DTO from directly priming DNA synthesis itself.

~~~
5’-CTTCATCCACGTTCACCTTGCCCCACAGGGCAGT<u>AACGGCAGACTTCTCCTC</u>/iSp9/
**CCCTATCCCTATCCCTA<u>ACACTAACCCAAACCCAA</u>/3’-Phos/**
~~~

The DTO contains exactly one binding site for the telg primer (underlined in the above text) and exactly one binding site for the HBBd primer (also underlined). After extending DNA synthesis from the 3’ end of the hybridized telg primer along the DTO, the DNA polymerase is blocked from further extension by the C9 linker. This feature ensures that telg extension along the DTO will not interfere with HBBd primer hybridization and extension. The bases depicted in red font are purposely mutated from the natural telomere sequence.

The order for the DTO was for 250 nmoles, PAGE-purified. The yield that was shipped was 11 nmoles of dried down DTO, which we dissolved in 10 mM Tris-Cl, 0.1 mM EDTA, pH 8.0 to a stock concentration of 11 uM.

### MMqPCR

PCR reactions are set up by aliquoting 5 ul of a 2x MasterMix containing all four primers into each reaction well, followed by either 5 ul of the subject’s purified DNA sample (approximately 3 ng of DNA) or, for the Standard Curve reactions, 5 ul of one of the serially diluted DTO concentrations, for a final volume of 10 ul per reaction. A dilution buffer, 10mM Tris–HCl, 0.1mM EDTA, pH 8.0, including 100 ng BSA (bovine serum albumin) per ul, is used to dilute concentrated stocks of the subjects’ DNA samples and the DTO to these much lower working concentrations for qPCR.

The composition of the 2x MasterMix is: 100 mM Tris, pH 9.2; 32 mM ammonium sulfate; 0.1% Brij 58; 10 mM MgCl_2_; 2M betaine; 400 nM of each dNTP; 2x SYBR Green I; 2x Titanium Taq DNA Polymerase (Takara Bio.); 100 ng BSA per ul; telg primer, 300 nM; telc primer, 400 nM; HBBu primer, 1 uM; and HBBd primer, 1 uM. This mix is routinely prepared beginning with the RB20 10x reaction buffer from DNA Polymerase Technology, Inc., and then adding betaine, dNTPs, SYBR Green, DNA polymerase, BSA, primers, and additional MgCl_2_.

All experimental DNA samples are assayed in triplicate. Standard Curve reactions were assayed in quadruplicate.

The high end concentration of the Standard Curve serial dilution is prepared by diluting the stock 11 uM DTO solution with the above-mentioned dilution buffer, to a concentration of approximately 3.9 × 10^6^ copies per ul. Eleven additional 3-fold serial dilutions from this high end concentration are then prepared, down to the lowest concentration of approximately 22 copies per ul. The middle two concentrations from this set of 12 DTO concentrations are then discarded. The remaining highest 5 DTO concentrations are used for the telomere Standard Curve, and the remaining lowest 5 DTO concentrations are used for the HBB Standard Curve.

Reactions are carried out in a Hard-Shell white wall 384-well plate compatible with the Bio-Rad CFX384 Real-Time PCR Detection System.

Exponential amplification from the telomere and single copy gene (HBB) targets with SYBR Green I fluorescent signal acquisition in a monochrome multiplex closed tube assay is then performed:

**Figure 1.**
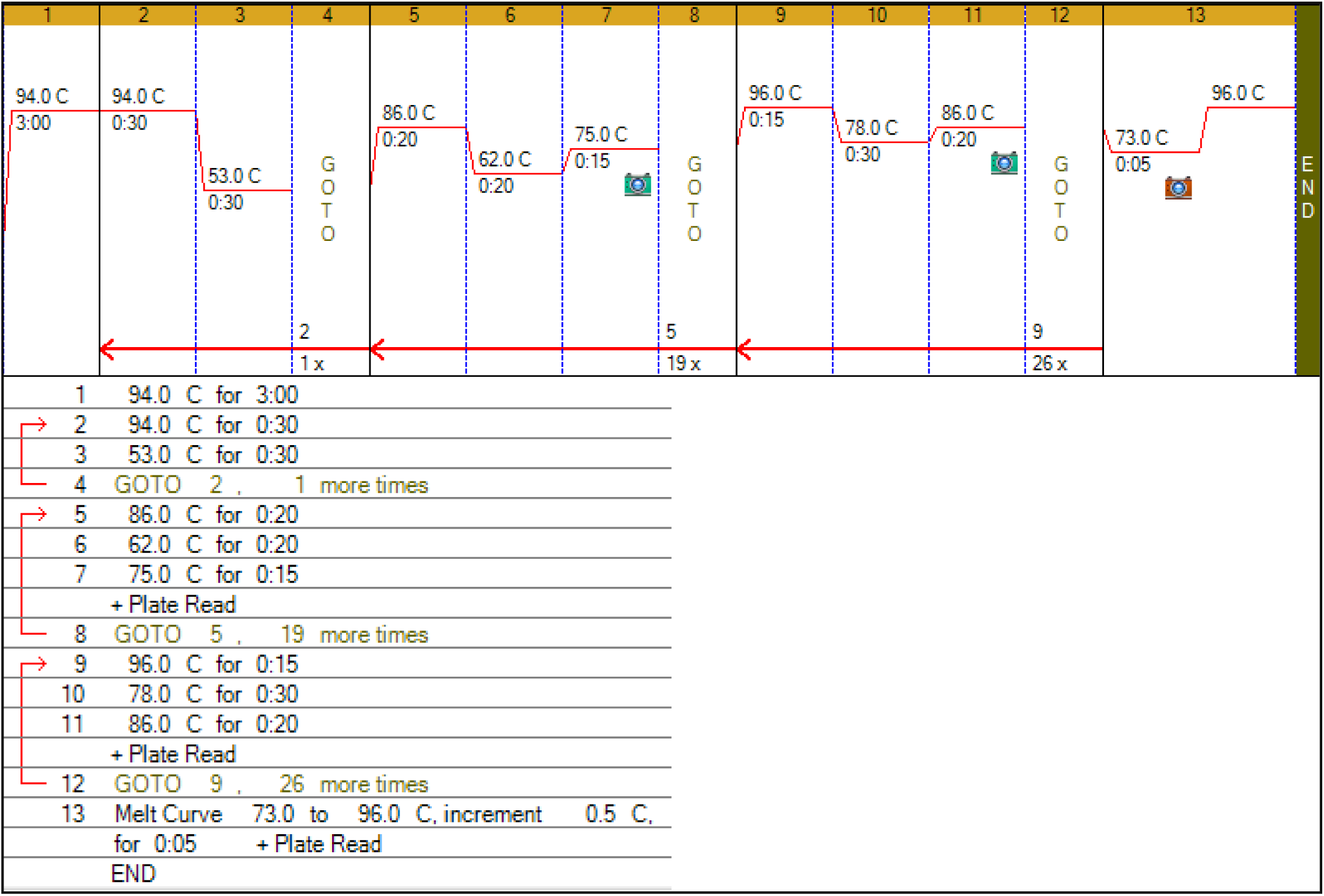
Thermal profile.

Steps 2 and 3, × 2 cycles, accomplishes the initial priming and copying of both telomere and HBB sequences. The telomere amplification signal is then collected at 75.0°C over 20 cycles in which the highest temperature reached is 86°C, sufficient to melt the telomere amplicon, but too low to melt the GC-clamped HBB amplicon, thereby putting its amplification on hold. The following stage consists of 27 cycles of high temperature thermal cycling when the HBB target is exponentially amplified, with signal acquired at 86.0°C sans interference from the telomere products synthesized earlier, which remain fully melted and therefore unable to affect the fluorescent signal during this stage.

After thermal cycling and raw data collection are completed, the Bio-Rad CFX Manager 3.1 software is used to generate two standard curves for each plate, one for the telomere signal (collected at Step 7), using the 5 highest DTO dilutions, and one for the HBB signal (collected at Step 11), using the 5 lowest DTO dilutions.

The T value for each DNA sample is the Standard Curve DTO dilution that contains the same number of copies of the telomere sequence as the experimental sample, and the S value is the DTO dilution that contains the same number of copies of the single copy gene sequence as the experimental sample. Dividing the first dilution by the second dilution yields an absolute T/S ratio, since it is expressed relative to the fixed 1:1 T/S ratio that is built into the DTO by design. T/S is proportional to the average telomere length per cell in the tissue sample from which the genomic DNA was extracted.

## RESULTS

### Independent standard curves for telomere and HBB

**Figure 2.**
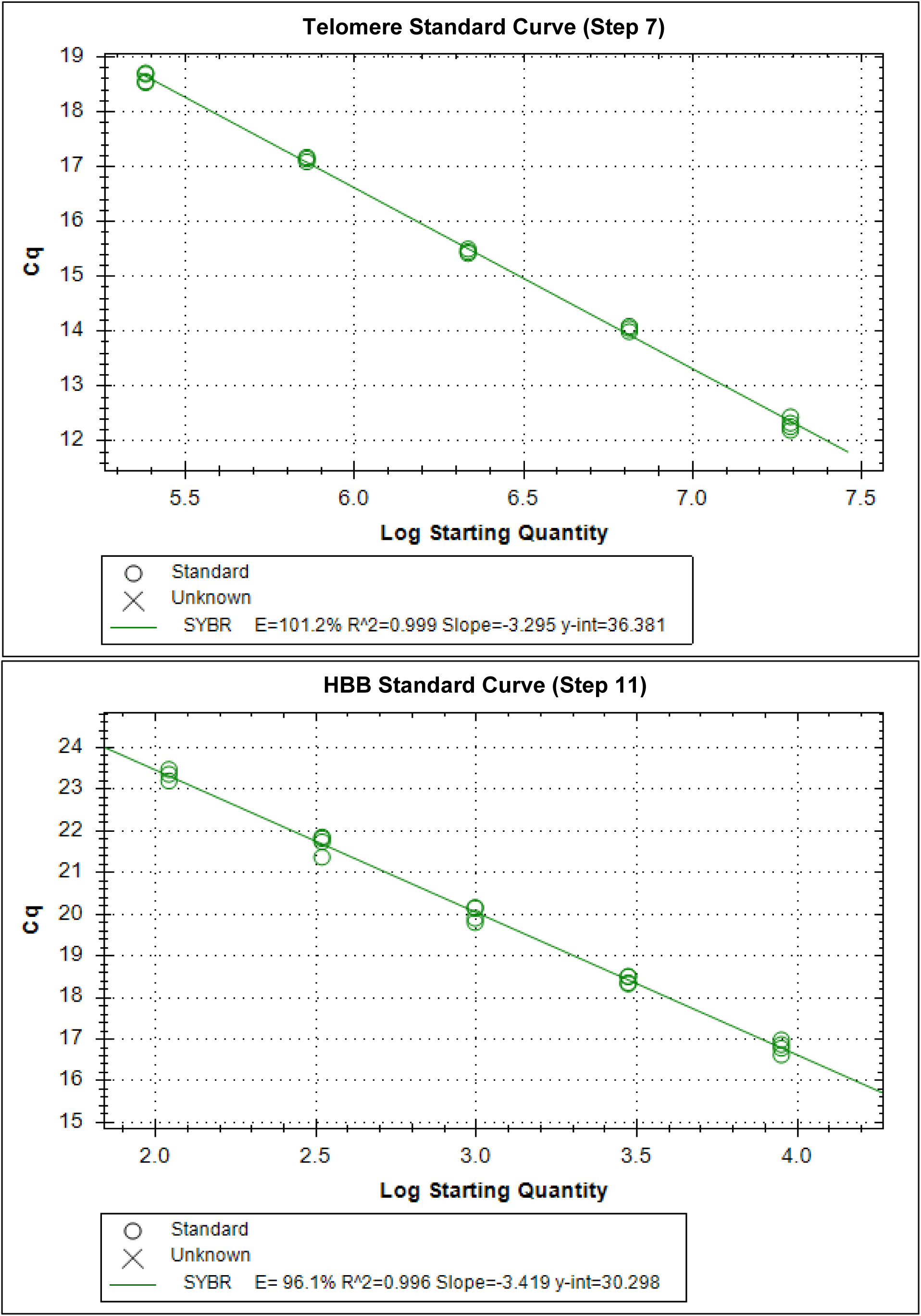
Standard Curves. The Starting Quantity is the copy number of the DTO.

### Melting profiles of telomere and HBB PCR products

**Figure 3.**
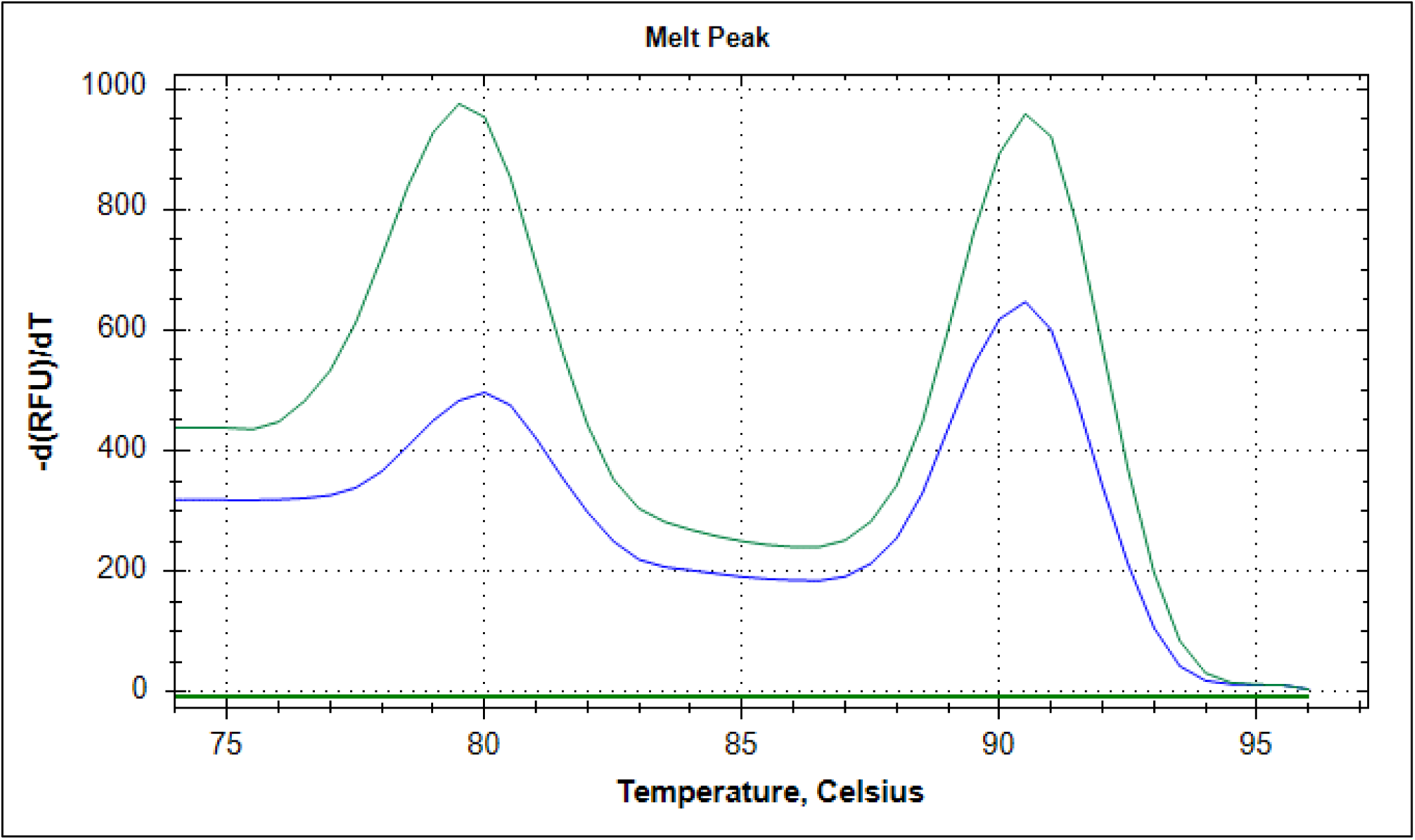
Melting profiles (dissociation curves) for the telomere and beta-globin (HBB) products, obtained at the end of the MMqPCR. Green curve: starting template at the beginning of the MMqPCR was genomic DNA; blue curve: starting template was the single-stranded dual-template oligonucleotide (DTO). The peak at about 80°C is for the telomere PCR product, and the peak at about 90.5°C is for the GC-clamped HBB PCR product.

### Correlation between mTRF lengths and T/S ratios

**Figure 4.**
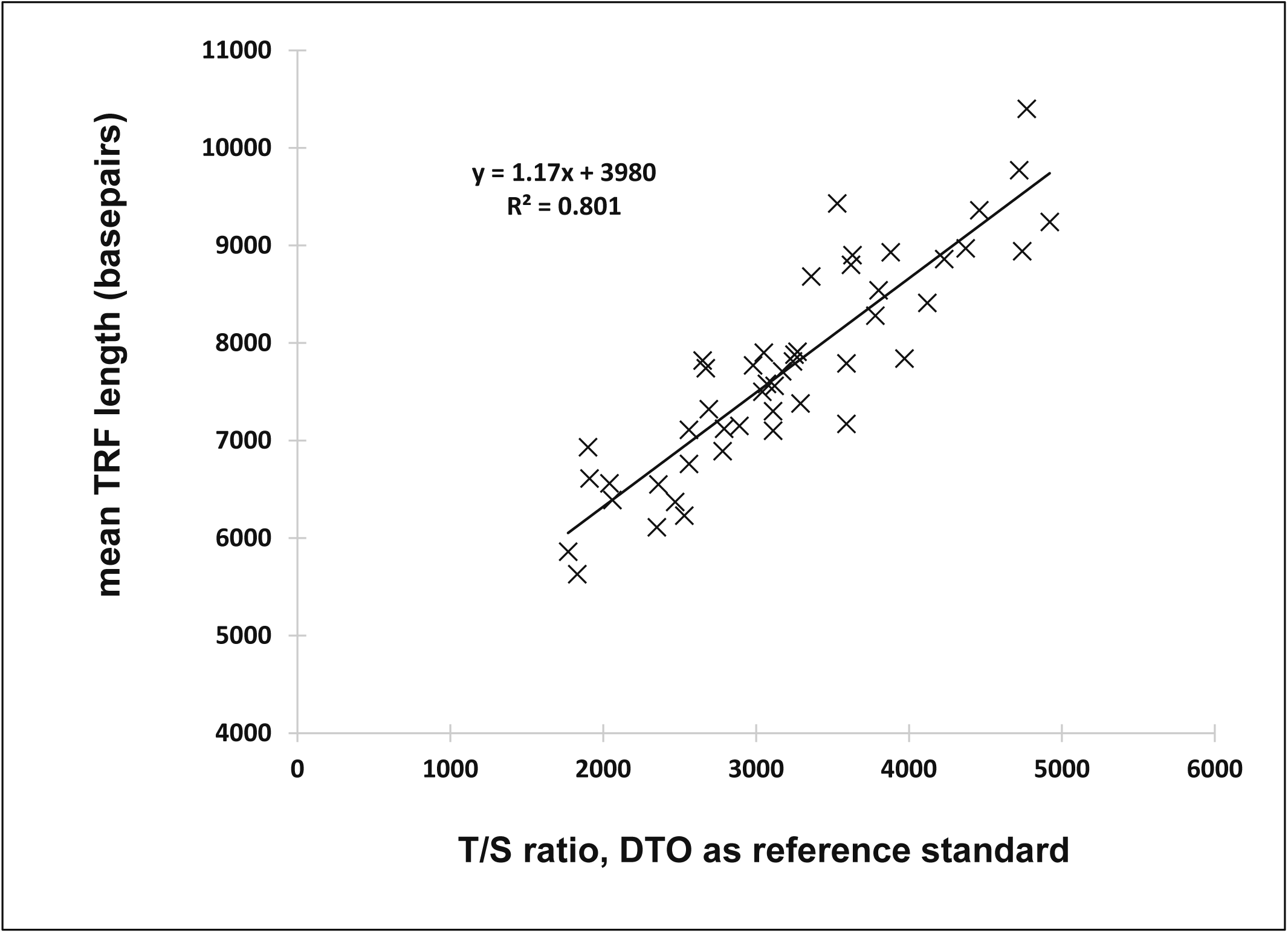
T/S ratios and mean Terminal Restriction Fragment (TRF) lengths for 48 Utah CEPH individuals. The y-intercept (3980 bp) is interpreted as the mean length of the subtelomeric DNA segment between the restriction enzyme’s cut site and the beginning of the true telomere sequence repeats. The intra-assay geometric mean of the coefficient of variation of T/S for these 48 DNA samples was 5.99%.

### Decline in T/S with aging

**Figure 5.**
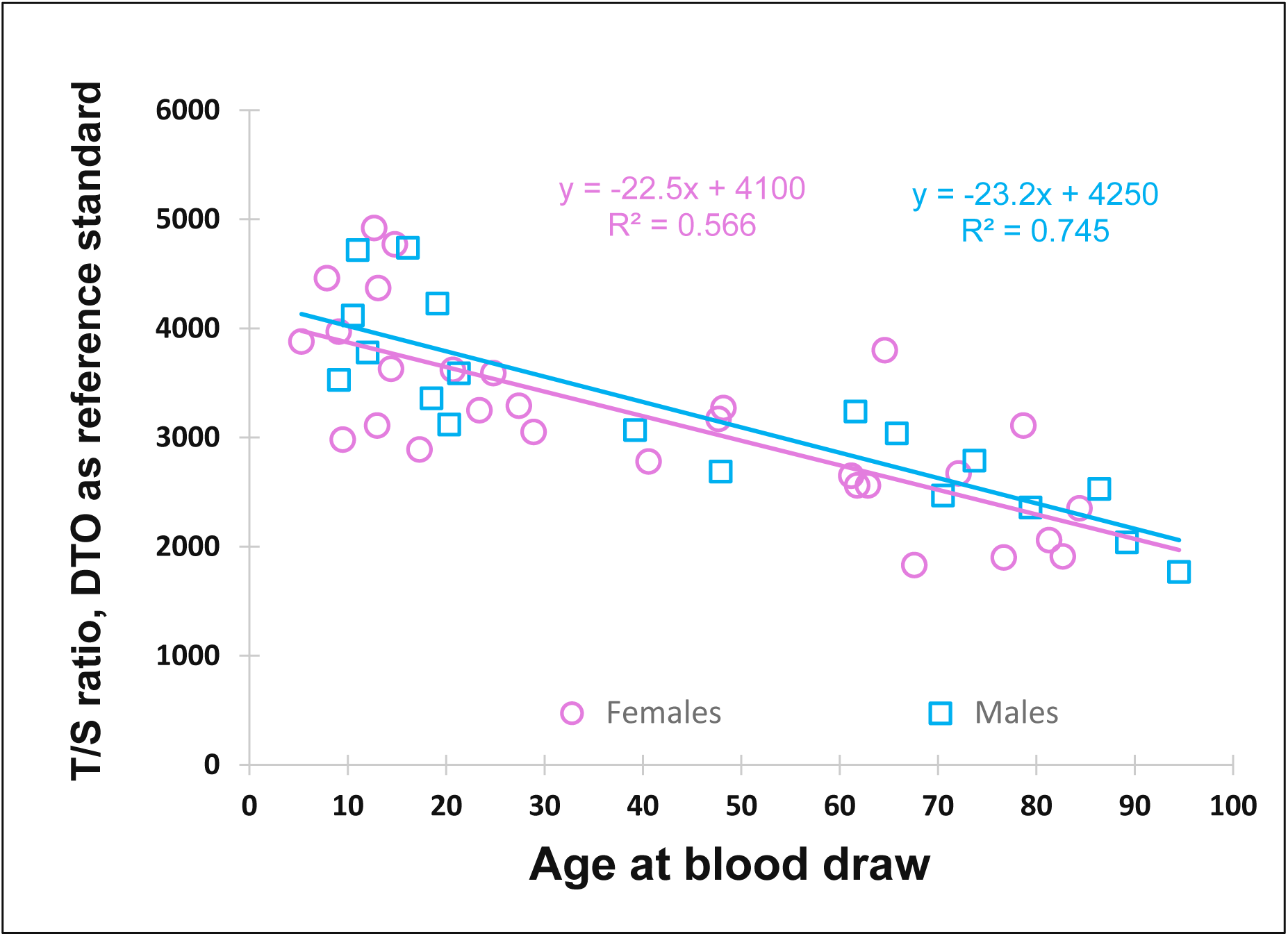
T/S ratios decline with age in both females and males. Pink circles, females (n=29); blue squares, males (n=19).

## DISCUSSION

Here we have shown that a single-stranded dual-template oligonucleotide (DTO) can be used as the reference standard for measurements of average telomere length by quantitative PCR. The T/S result for a given experimental DNA sample is a ratio of two dilutions of the DTO: the dilution that matches the DNA sample on copy number of the telomere sequence, divided by the dilution that matches the DNA sample on copy number of the single copy gene (scg). This can be considered an absolute T/S ratio, as it is not expressed relative to the T/S ratio of some natural genomic DNA sample, but rather is expressed relative to the known 1:1 telomere:scg ratio of the DTO. However, whether the T/S values for a set of DNA samples obtained using one batch of the DTO will be faithfully preserved when the assay is repeated using an independently prepared batch of the DTO has not yet been tested.

In principle these absolute ratios can be converted to approximate average telomere lengths in basepairs of DNA, by simply multiplying the slope of the regression line in Figure 4 (where slope = 1.17) by the T/S value of each experimental DNA sample. However, whether that slope value will be the same across various cohorts and human populations has not yet been tested. The use of more exact experimental approaches to derive the most accurate factor to use when converting these DTO-based absolute ratios to telomere lengths in basepairs is beyond the scope of the current study.

In developing this assay, we used an ammonium sulfate based PCR buffer with no potassium chloride (KCl), instead of more typical PCR buffers that contain KCl, because the G-rich telomeric strand is, in principle, prone to forming G-quadruplex structures (G4s) in the presence of KCl, and G4s elsewhere in the genome have already been reported to sometimes interfere with PCR when KCl-containing buffers are used, but not when ammonium sulfate buffers lacking KCl are used [7,8].

We chose Titanium Taq as the DNA polymerase because it is compatible with a wider range of MgCl_2_ concentrations than standard Taq polymerase, thereby providing more flexibility in optimizing PCR reactions; and also because it is more thermostable than standard Taq polymerase.

This new assay is simple, rapid, and readily scalable to achieve a high throughput of samples. The range and decline of T/S values across the human lifespan obtained using this assay are consistent with previously reported values. We hope that this DTO and the accompanying protocol will facilitate the standardization of average telomere length measurements and analyses across laboratories, and prove useful in the investigation of the biology of telomeres and the roles they play in the molecular pathophysiology of multiple diseases and aging.

## ACKNOWLEDGEMENTS

We thank all Utah individuals who participated in the CEPH consortium. We thank Mark F. Leppert for access to the Utah CEPH DNA samples.

## FUNDING

This work was supported by NIH R21 AG054962 to Dr. Cawthon.

## COMPETING INTERESTS

Dr. Cawthon and the University of Utah have licensed the earlier versions of our patented qPCR methods for measuring average telomere length (references 5 and 6) to Telomere Diagnostics, Meno Park, CA.

